# Calcium-modulated *cis* and *trans* E-cadherin EC1-2 interactions play a key role in formation, dynamics and plasticity of cadherin junctions

**DOI:** 10.1101/2024.10.14.618140

**Authors:** J. Lai-Kee-Him, A. Fouillen, M. Chee, N. Lemercier, H. Feracci, P. Bron

**Author notes:** Imagxcell, 2 allée des Feuillantines 94800 Villejuif (N. Lemercier). Correspondence: Patrick Bron, Centre de Biochimie Structurale, INSERM U1054 / CNRS UMR 5048 and Université Montpellier, France.

## Abstract

Classical cadherins are ubiquitous calcium-dependent adhesive glycoproteins required in junction formation in metazoans. Calcium ions are essential for cadherin biological activity, but how they act in synergy is still a matter of controversy. We investigated by cryo-electron microscopy and cryo-electron tomography, a biomimetic system consisting of liposomes decorated with the two first E-cadherin extracellular modules, previously reported as functionally essential in junction formation. We showed that E-cadherin extracellular modules 1 and 2 are sufficient to form junctions in presence of calcium. Together with the first adhesive contacts, we observed the formation of cadherin patches and quasi-crystalline structures on the free surface of proteoliposomes suggesting that calcium complexation induces the self-interaction of the E-cadherin extracellular modules 1 and 2 through *cis* interactions. We propose that these calcium-dependent cadherin nanoplatforms, by facilitating the formation frequency of *trans* interactions, most probably favor the formation of junctions. We also observed a cooperative effect between *trans* and *cis* interactions leading to the protein depletion of liposome surfaces. Additionally, Cryo-electron tomography experiments performed on extracellular E-cadherin modules 1 and 2-mediated junctions revealed the coexistence of different molecular arrangements previously described from crystallography studies. Thus, this study provides a new insight into biophysical mechanisms underlying how calcium binding modulates cadherin adhesive function, in showing that E-cadherin 1 and 2 modules play a key role through their intrinsic ability to form multiple calcium-dependent *cis* and *trans* interactions, in the formation of cadherin junctions contributing to the dynamics and plasticity of cadherin-mediated cell junctions.

## INTRODUCTION

Cell to cell adhesive contacts are specialized regions of the plasma membrane with unique morphologies and biochemical compositions that play essential roles in several cellular processes including morphogenesis, differentiation, proliferation and migration. Deregulation in cell adhesion can lead to numerous pathologies such as inflammation, the invasion of a tissue by a pathogen, the body’s reaction to biomaterials, and in particular to the formation of metastases (Jeanes, Gottardi, & Yap, 2008). Proteins involved in cell adhesion can be classified into 4 structural families: proteins related to (a) immunoglobulins, (b) integrins, (c) selectins and (d) cadherins (Buckley, Rainger, Bradfield, Nash, & Simmons, 1998). The latter form one of the main families of molecules that are responsible for Ca^2+^-dependent homophilic adhesion between cells (Kemler, 1992; Maître & Heisenberg, 2013) and is composed of four major subfamilies: classical cadherins, desmosomal cadherins, protocadherins and atypical cadherins.

Classical cadherins are single-pass transmembrane glycoproteins with an extracellular segment consisting of five similar tandemly arranged cadherin domains (EC) named EC1-EC5. Each domain has an Immunoglobulin-like structure consisting of seven antiparallel ß-strands organized into two opposing sheets (L. Shapiro et al., 1995) and are relied by linkers, each comprising three calcium binding regions (Brasch, Harrison, Honig, & Shapiro, 2012). Starting from the N-terminal end (Nollet, Kools, & van Roy, 2000), the EC1-5 segment is followed by a transmembrane region and a highly conserved cytoplasmic domain. The intracellular signaling pathway initiated by cadherins is due to an association of the intracellular domain with the actin cytoskeleton via cytoplasmic proteins such as catenins, plakoglobin and p-120 (Delmas et al., 1999; Ozawa, Baribault, & Kemler, 1989; Yap, Brieher, & Gumbiner, 1997).

Among classical cadherins, we find epithelial (E)-, neuronal (N)-, and vascular endothelial (VE)-cadherins that form adhesion between epithelial, neuronal and vascular endothelial cells, respectively. Classical cadherins can be subdivided into type I and type II. Type I classical cadherins, comprising E- and N-cadherins, have conserved *(a)* HAV tripeptide motif (Blaschuk, Sullivan, David, & Pouliot, 1990) and *(b)* tryptophan 2 (Trp 2) residue (L. Shapiro et al., 1995) in the most distal EC (EC1). Type II classical cadherins, as VE-cadherins comprise two conserved Trp (Trp 2 and Trp 4) (Lawrence Shapiro & Weis, 2009), while the HAV motif is absent.

E-cadherin is mainly localized at the adherens junctions in the basolateral membrane of polarized epithelial cells. It is also involved in embryonic development (Geiger & Ayalon, 1992), or later during development and in adult tissues, where it helps maintaining epithelial cell polarization and tissue integrity (Larue et al., 1996; Yap, Brieher, Pruschy, & Gumbiner, 1997). The adhesive function of E-cadherin is frequently lost during the development of most, if not all, human epithelial cancers (Perl, Wilgenbus, Dahl, Semb, & Christofori, 1998). Thus, E-cadherin is central to epithelial biogenesis and stabilization.

E-cadherin comprises four conserved binding pockets (PENE (residues 10-13), LDRE (residues 66-69), DQNDN (residues 100-104), and DAD (residues 134-136)) positioned between successive EC modules that bind each three Ca^2+^ ions in a highly cooperative manner (Nagar, Overduin, Ikura, & Rini, 1996). Some single point substitution of some motifs, as for example the substitution of the Asp103 (D103) to Lys/Ala from the DQNDN motif, result in markedly reduced adhesion (Handschuh, Luber, Hutzler, Höfler, & Becker, 2001). The cadherin ectodomain is fully Ca^2+^-occupied under physiological conditions, as the Ca^2+^ concentration in the extracellular media (approximately 1 mM) (Sotomayor & Schulten, 2008) is higher than binding affinities of the Ca^2+^ sites (average dissociation constant K_D_ ≈ 25 µM). Subsequent to this Ca^2+^ binding, the cadherin ectodomain is believed to adopt a rigid rod-like conformation that will favor *cis*- or *trans*-interaction (Cailliez & Lavery, 2005; L. Shapiro et al., 1995), it means interactions between E-cadherins located on the same cell surface (*cis*) or from two opposite cells (*trans*).

Structural studies performed on cadherins are tremendous, mainly revealing that classical cadherin ectodomains can interact through the formation of a strand-swap dimers (S-dimer) or X-dimers. A complete and well documented overview of the current events involved in the formation of E-cadherin adhesions, are proposed in (Biswas & Zaidel-Bar, 2017; Priest, Shafraz, & Sivasankar, 2017). The S-dimer consists in swapping the N-terminal β-strands of their EC1 modules, with docking of the highly conserved Trp 2’s lateral chain into the hydrophobic pocket of the paired molecule, and salt bridge interactions formed by positively charged N-terminal residues. The X-dimer results in extensive surface interactions along the EC1-2 interdomain linker region. It was shown by nuclear magnetic resonance (NMR) that X-dimers initiate cadherin dimerization followed by switching to S-dimer (Li et al., 2013), but also that E-cadherin ectodomains can shuttle between X-dimer and S-dimer conformations (Manibog et al., 2016) indicating that the X-dimer is not a transient intermediate state in the formation of the strand-swap dimers (Harrison et al., 2010; Sanjeevi Sivasankar, Zhang, Nelson, & Chu, 2009).

For a long time, the admitted general model explaining the formation of E-cadherin junctions proposed that *trans* adhesive calcium-dependent cadherin interactions from two opposite cells were first established (Perez & Nelson, 2004) followed by *cis* interactions allowing the recruitment of additional E-cadherins (Wu et al., 2010; Wu, Vendome, Shapiro, Ben-Shaul, & Honig, 2011). The interplay between *trans* and *cis* interactions together with a complex molecular scaffold on the cytoplasmic side, must participate in the formation and dynamics of cellular junctions. Nonetheless, the abundant literature on studies performed with various experimental approaches, and dealing with cadherins in solution or linked onto surfaces, as well as with cadherin-expressing cells, has raised some inconsistencies. For example, Takeda et *al.* (Takeda, Shimoyama, Nagafuchi, & Hirohashi, 1999) used chemical cross-linking studies on cells to show that Ca^2+^ ions induce lateral *cis* E-cadherin dimer formation at the cell surface, supporting the formation of Ca^2+^-dependent *cis* interactions. In line with this, molecular dynamic studies indicate that calcium ions favor the stabilization of a *cis*-dimer complex formed between EC1-2 fragments (Cailliez & Lavery, 2005). The role of *cis* E-cadherin interactions outside the context of trans-interaction was suggested but not unambiguously established. Moreover, it was also proposed that the molecular mobility of E-cadherins plays a role on the assembly of cadherins adhesions, and consequently the importance of cadherin interaction with actin and spectrin cytoskeleton underlying the cell membrane (Engl, Arasi, Yap, Thiery, & Viasnoff, 2014; Erami, Timpson, Yao, Zaidel-Bar, & Anderson, 2015). Indeed, it was shown that immobile fraction of E-cadherin molecules serve as the seed for nucleating extended cadherin adhesions (Biswas et al., 2015). These observations have to be related with quantitative Forster resonance energy transfer (FRET) approaches demonstrated that E-cadherin forms constitutive dimers in plasma membrane and that these dimers exist independently of the actin cytoskeleton or cytoplasmic proteins (Singh, Ahmed, Sarabipour, & Hristova, 2017). More recently, with the development of high-resolution microscopies, it was shown that cadherin extracellular domain can form clustering in the absence of *trans*-interactions using mimetic systems, resulting from a combination of specific and nonspecific *cis*-interactions (Thompson et al., 2020; Thompson, Vu, Leckband, & Schwartz, 2019), and that cadherin *cis* and *trans* interactions were shown as mutually cooperative (Thompson, Vu, Leckband, & Schwartz, 2021). These data indicate that the sequence of events leading to the establishment of cadherin-mediated junctions, and in particular the *trans* vs *cis* interaction has not been conclusively elucidated yet. Others important considerations come from electron microscopy (EM) studies on adhesive structures such as *adherens junctions* (Tariq et al., 2015) and desmosomes using tomography (He, Cowin, & Stokes, 2003) which suggest various putative molecular interfaces *in situ* for *cis* and *trans* interactions. Consequently, it is still compulsory to improve the understanding of biophysical mechanisms underlying how calcium binding modulates cadherin adhesive function, *cis*- and *trans*-dimer equilibrium and organization of cadherins in adhesive clusters.

In this context, we used cryo-EM and cryo-electron tomography (cryo-ET) to investigate how calcium ions impact the adhesive homophilic interactions between E-cadherin fragments linked to liposomes. We decided to focus on the pair modules EC1-2 from the E-cadherin (E/EC1-2), as this distal segment is essential for E-cadherin biological activity and contains the integrity of the outermost binding pocket necessary for the proper adhesive activity of the full-length molecule. Indeed, it has been shown that the E/EC1 module governs the first steps of adhesion, its efficiency and its specificity (Shan, Koch, Murray, Colman, & Shapiro, 1999) and that these pair modules are essential i) for cadherin-mediated adhesion between cells (Gumbiner, 1996), ii) between surfaces covered with these fragments (Leckband & Sivasankar, 2012; Emilie Perret et al., 2002), and iii) between cells and decorated surfaces (Arulanandam et al., 2009; Chevalier et al., 2010). We also used a single point mutation of the D103 to Ala (E/D103A) of the calcium-binding pocket of the E/EC1-2 fragment. This E/D103A substitution on the full-length E-cadherin is reported to have a negative effect on cadherin-mediated cell-cell adhesion (Handschuh et al., 2001) whereas in E/EC1-2 fragment, the mutation limits Ca^2+^ binding and increases the relative mobility of EC1 and EC2 modules even though the ß-sheet folding characteristics of cadherin modules are preserved.

## RESULTS

### EC1-2 decorated liposomes form calcium-dependent adhesive contacts

First, calibrated liposomes of about 100 nm in diameter, designed from synthetic DOPG (1,2-dioleyl-sn-glycero-3-phospho-(1’-rac-glycerol) (sodium salt)) and DGS (1,2-dioleoyl-sn-glycero-3-succinate) nickel-nitrilotriacetic acid (Ni-NTA) functionalized lipids, were prepared and either quickly frozen or after ten minutes incubating time in liquid ethane for cryo-EM observation. When observed by cryo-EM whatever freshly frozen or after up to one-hour incubation, they appeared as smooth spheres embedded into a thin layer of amorphous ice (Figure 1A). They were delineated by two parallel dark lines revealing the bilayer organization of the lipid membrane (see inset Figure 1A) and remained isolated without or with calcium into the buffer (Figure 1, compare A and B). We then decorated the NTA-liposomes with E/EC1-2 fragments (Figure 1C-E), in order to use them as a model system to decipher the formation of cadherin-mediated adhesive contacts. Their ability to aggregate was examined depending on calcium conditions and E/D103A-decorated liposomes were used as a putative non-adhesive control. In order to be under physiological conditions, the calcium concentration was fixed to 1 mM. All recombinant cadherin fragments used here were expressed with a C-terminal hexahistidine tag, allowing their oriented binding to liposomes thanks to the Ni-NTA moiety of the DGS functionalized lipids. Following incubation with E/EC1-2 fragments, liposomes exhibited a uniform coverage of protruding densities (Figure 1C, black arrows) reflecting the binding of cadherin fragments through their C-terminal hexahistidine tag to Ni-NTA DOGS lipids, uniformly dispersed at the surface of the lipid bilayer. As expected, without calcium, E/EC1-2-liposomes remained scattered (Figure 1C) while, after a few minutes in the presence of 1 mM calcium, they started to display numerous contact areas that we henceforth refer as junctions (Figure 1D). These junctions were thus brought about by calcium-dependent *trans* adhesive interactions between E/EC1-2 fragments positioned on facing liposomes. As pointed out by black arrows in Figure 1D, these adhesive contacts were able to induce local flatness of liposomes membranes, suggesting that the collective effect of E/EC1-2 intermolecular interactions must be quite robust. In contrast, liposomes decorated with E/D103A fragments remained scattered in solution even in presence of calcium (Figure 1E).

**Figure 1 :**
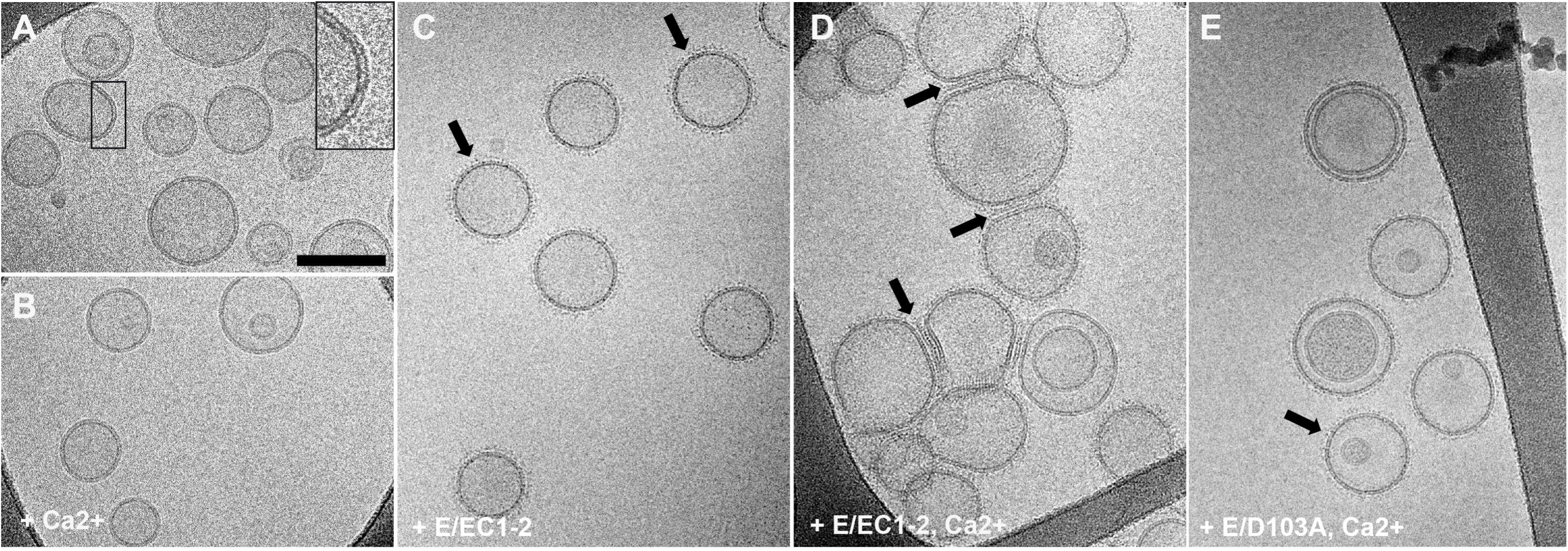
Cryo-EM observations of liposomes and EC1-2 decorated liposomes. **A and B** : Liposomes made from DOPG/DOGS-Ni-NTA (4/1 : w/w) lipid mixture (**A**) in absence or (**B**) in presence of 1 mM calcium (Ca^2+^) after 10 min incubation. **C and D** : E/EC1-2 decorated liposomes (**C**) in absence or (**D**) in presence of 1 mM calcium (Ca^2+^) after 10 min incubation. The interplay of E/EC1-2 fragments with liposomes is reflected by the presence of black densities protruding out the surface of liposomes (black arrows). (**C**) While isolated without calcium, (**D**) E/EC1-2 decorated liposomes form numerous junctions after calcium adding. **E** : E/D103A decorated liposomes incubated 10 minutes with 1 mM calcium (Ca^2+^). Scale bar, 100 nm.

### Calcium ions favor E/EC1-2 lateral clustering

In an attempt to explore the sequence of events leading to E/EC1-2 junction formation, we examined more closely the behavior of E/EC1-2 decorated liposomes upon 1 mM calcium addition. We designed our experiments with cryo-EM observations of E/EC1-2 decorated liposomes after 5, 10, 15 and 30 min calcium incubation. Figure 2 shows representative views of E/EC1-2 junctions observed between 5 and 15 min of incubation with calcium.

**Figure 2 :**
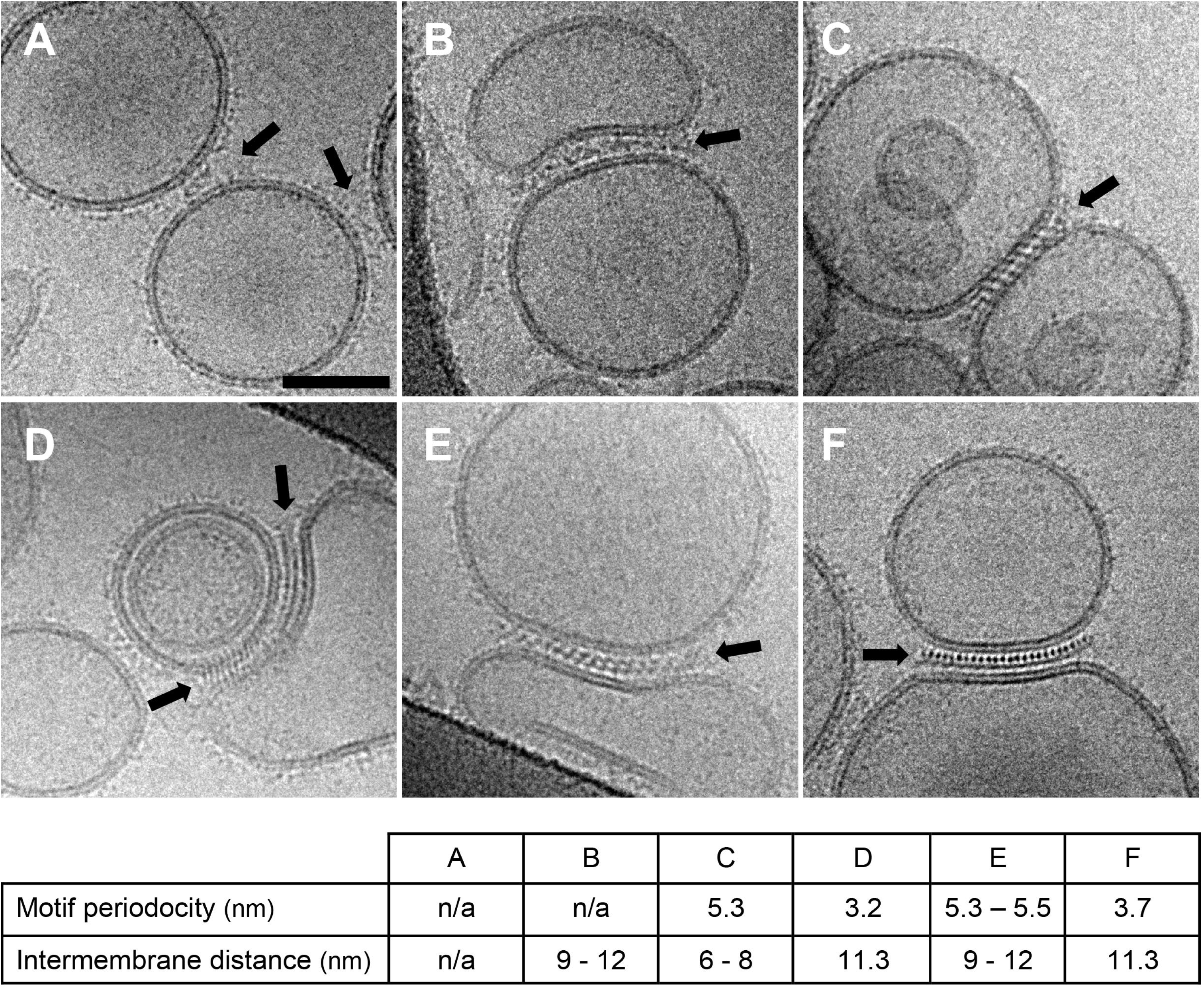
Representative views of E/EC1-2 junctions. **A**: Representative view of a junction formed by the interplay of a few E/EC1-2 fragments (black arrows). **B** : Representative view of non-organized junction (black arrow). **C – F** : Various views observed with well-structured E/EC1-2 junctions (black arrows). The table reports the motif periodicity and inter-membrane distance in nm computed from A to F inserts. Scale bar, 50 nm.

As expected and observed in Figure 1C, the main visible effect of calcium ions was the formation of well-ordered contact areas between liposomes (Figure 2), that were visible for a calcium incubation time up to 15 min but never at and after 30 min (see below). Three types of junctions could be observed: small adhesive structures involving only a few E/EC1-2 fragments (Figure 2A), larger and disordered junctions (Figure 2B), and well-structured contacts displaying various repetitive patterns (Figure 2C-F). The most unexpected observation was the presence of numerous E/EC1-2 semi-crystalline two-dimensional (2D) arrays on free proteoliposomes’s surface (Figure 3), exclusively observed in the range of 5 to 10 min calcium incubation. These 2D arrays were either observed within proteoliposomes involved in junctions or within isolated proteoliposomes (Figure 3A versus 3F). They displayed discrete peaks adopting a rectangular pattern in the power spectrum of their Fourier transform (FT) (Figure 3B and insets in D, E and F). The presence of regular peaks in FT of images indicates that E/EC1-2 fragments adopted a regular organization at the surface of liposomes. But only one order of diffraction is observed, suggesting more a semi-crystalline organization for E/EC1-2 fragments.

**Figure 3 :**
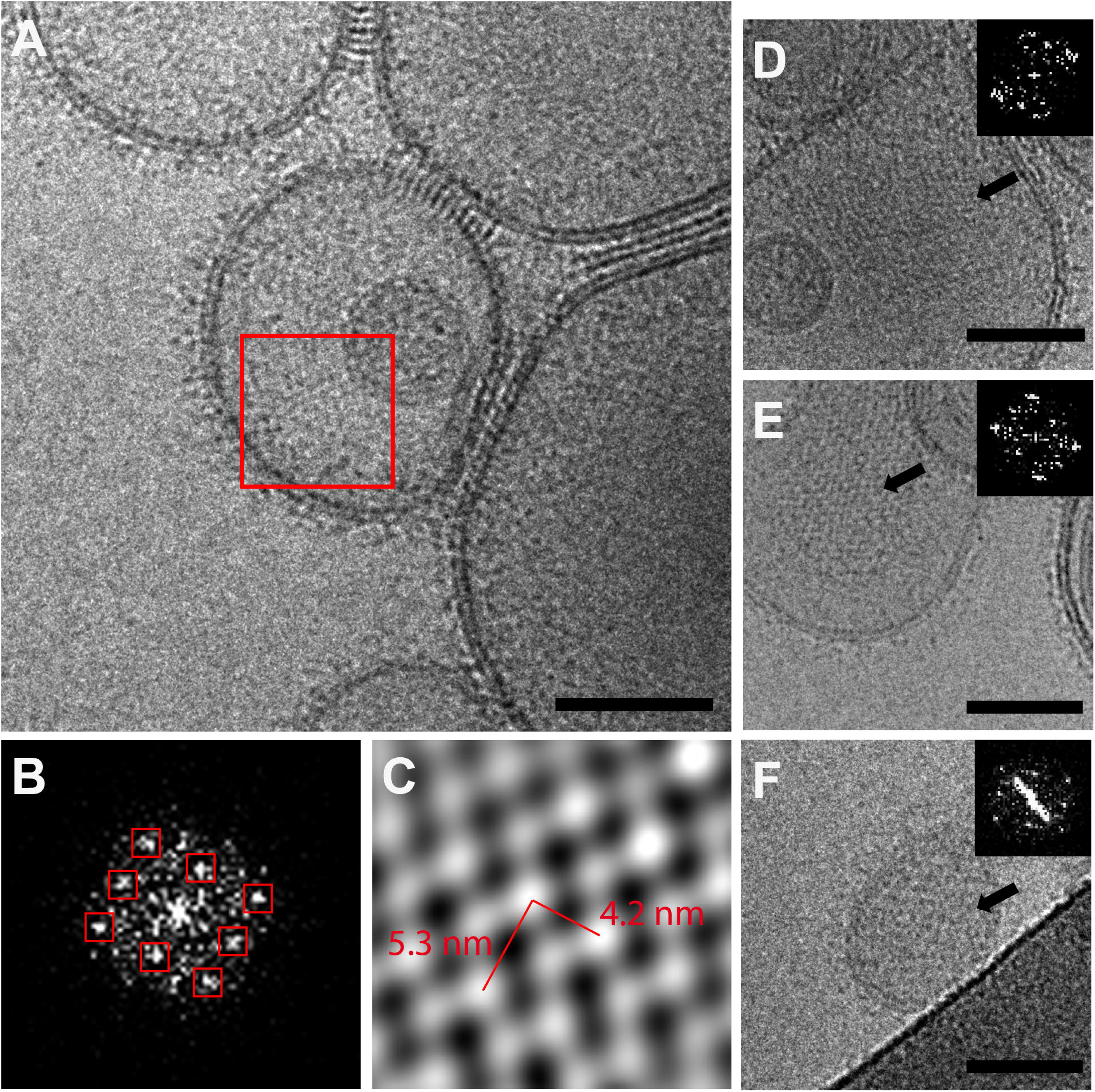
Lateral interplays between EC1-2 fragments. **A:** E/EC1-2 proteoliposomes after 10 min calcium (Ca^2+^) incubation at 1 mM. The red square outlines a regular organization of E/EC1-2 fragments at the surface of the liposome. **B:** The power spectrum (PS) of the Fourier transform (FT) of the red zone delineated in A. This PS shows regular peaks, squared in red, revealing the ordered nature of E/EC1-2 fragments. **C:** Filtered image obtained from the masked inversed Fourier transform presented in B. Here the signal used to compute the inversed FT is delineated by red squares in B. E/EC1-2 associated signal is in white. **D to F** : Various E/EC1-2 arrays and their corresponding PS. All PS inserts reveal regularly spaced peak forming a rectangle-like motif. Scale bars : 50 nm.

By applying masks around discrete peaks present in the FT (the signal inside masks being conserved) and computing the inversed FT, it is possible to get a filtered image where the signal comes from only the repetitive contribution present in the initial image, *ie* the regular E/EC1-2 fragments. In using this method, the filtered image of the region delineated by the red square in Figure 3A is presented in Figure 3C. The areas selected for this processing are indicated by red squares in Figure 3B. The filtered image (Figure 3C) exhibits repetitive white round domains, with distances separating them ranging from 5.3 to 4.2 nm. As images formed by the electron microscope correspond to 2D projections of a three-dimensional sample, it is difficult and hazardous to extrapolate from such projections a structural organization for E/EC1-2 fragments. The only reliable data that we can extract from these images is that E/EC1-2 fragments assemble in a regular array at the surface of liposomes involving *cis*-interactions between fragments after calcium addition. The formation of these 2D arrangements requires that cadherin molecules are close enough from each other, which is here possible due to the capacity of the cadherin fragments to move freely in the plane of the membrane. Such *cis*-arrays have never been observed with E/D103A-liposomes with or without calcium.

During the first 5 minutes of calcium incubation and preceding the formation of E/EC1-2 2D arrays, we regularly observed the local formation of small E/EC1-2 clusters (Figure 4A) at surface of liposomes, suggesting an early *cis*-interaction. As previously shown, between 10 and 15 min of calcium incubation, we observed both E/EC1-2 2D arrays and well-ordered contact areas between liposomes (Figure 4B). After 15 min of calcium incubation, junctions get progressively disorganized (Figure 4C), no more *cis*-arrays are observed and E/EC1-2 fragments seem to be always associated in patches. Between 30 min to one-hour calcium incubation, E/EC1-2 proteoliposomes are aggregated, with a complete protein depletion of the liposome surfaces except at the level of junctions, where most of E/EC1-2 fragments are concentrated (Figure 4D).

**Figure 4 :**
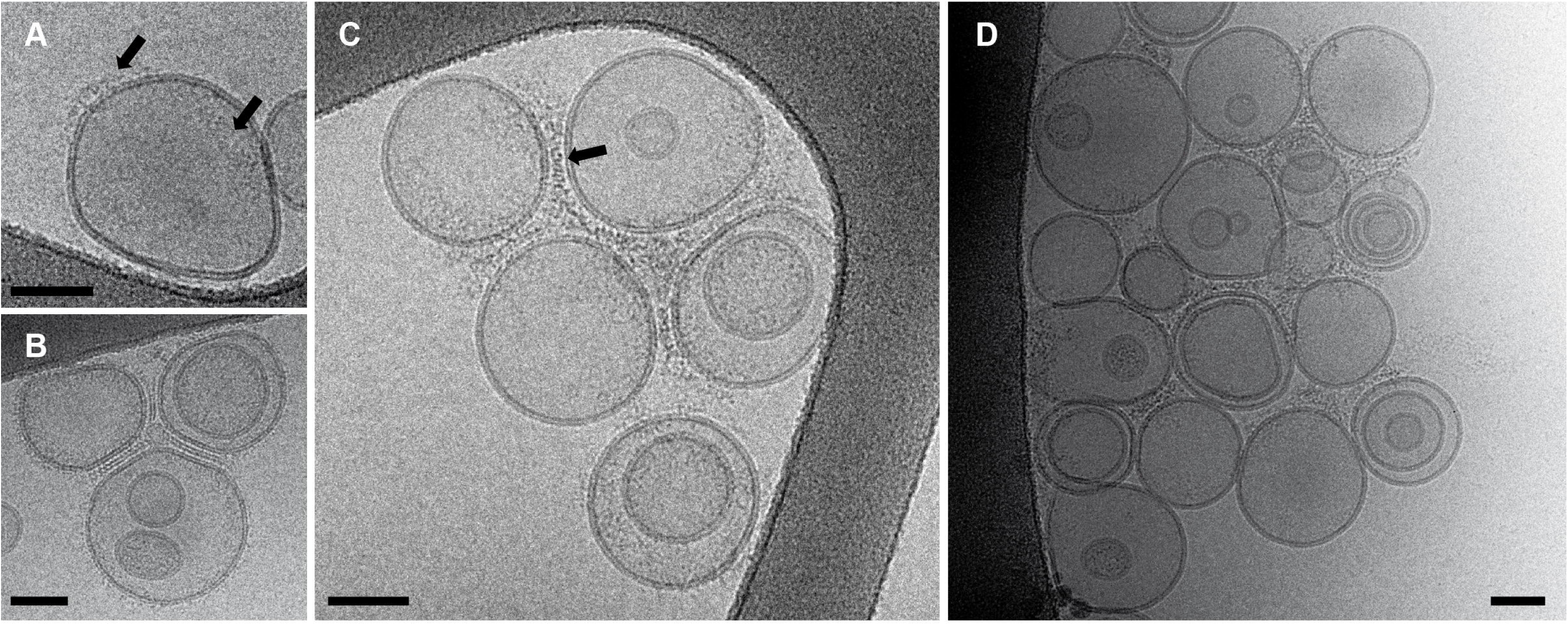
Chronological cryo-EM observations of E/EC1-2 decorated liposomes. **A:** E/EC1-2 proteoliposome observed after 5 min calcium (Ca^2+^) incubation at 1 mM showing earlier E/EC1-2 *cis* interactions inducing the fragments gathering **B:** E/EC1-2 proteoliposomes observed after 10 min calcium (Ca^2+^) incubation at 1 mM showing well-structured E/EC1-2 junctions **C:** Typical view of E/EC1-2 proteoliposomes observed after 20 - 30 min calcium (Ca^2+^) incubation at 1 mM showing progressive disorder of E/EC1-2 junctions. **D:** Typical view of E/EC1-2proteoliposomes observed after 30 min - 1 hour calcium (Ca^2+^) incubation at 1 mM showing complete protein depletion of the liposome surfaces. All E/EC1-2 fragments are localized at the level of junctions. Scale bars : 50 nm.

### Complexity of E/EC1-2-mediated *trans-*interactions

As previously observed, the most visible effect of calcium on E/EC1-2 decorated liposomes is the formation of well-ordered area contacts between liposomes (Figure 2), displaying various repetitive patterns (Figure 2C-F). The visible diversity of these patterns likely resulted from projections of regular arrays of E/EC1-2 fragments observed under different view angles. We then measured the inter-membrane distance between E/EC1-2 proteoliposomes forming junctions and the motif periodicity distances inside junctions (see table Figure 2). The motif periodicity distances observed are compatible with distances observed into E/EC1-2 2D-nanoplatforms whereas inter-membrane distances are mainly comprised between 9 to 12 nm. However, some junctions displayed shorter inter-membrane distances ranging from 6 to 8 nm. As E/EC1-2 length is about 9 nm from X-ray structures (Pertz, Bozic et al. 1999), the variation of inter-membrane distances suggested that junctions could adopt different E/EC1-2 molecular arrangements.

To go further in the understanding of the plasticity of E/EC1-2 junctions, we then used cryo-ET to study the organization of E/EC1-2-mediated junctions (Figure 5). The tilted-series images recorded from -60° to +60° are used to compute a tomogram, which is a 3D reconstruction of the sample. This provides the opportunity to move through the reconstructed volume and consequently through junctions. We can then explore the molecular architecture of E/EC1-2 modules within junctions and evaluate the minimal distance between membranes of two facing liposomes more accurately than from conventional cryo-EM images.

**Figure 5:**
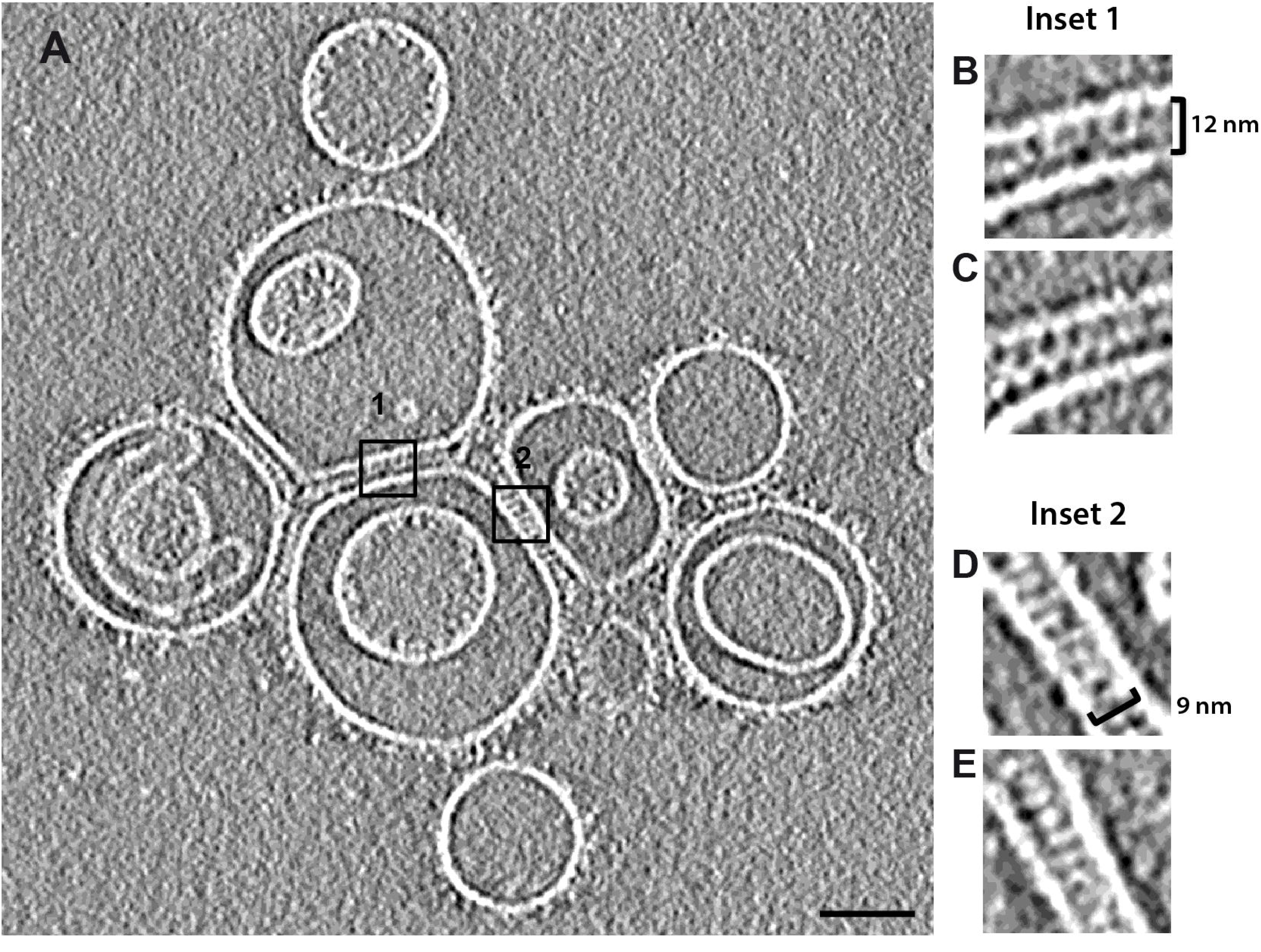
Cryo-electron tomography of E/EC1-2 junctions. **A** : cross section through the 3D tomogram. This section corresponds approximately to the central section, corresponding to section where the inter-membrane distances of junctions denoted 1 and 2 are minimal. The white continuous and linear densities correspond to lipid membrane while white densities between membranes correspond to EC1-2 fragments. **B and C** : Zoom in junction 1 for two different sections of the tomogram. **D** and **E** : Zoom in junction 2 for two different sections of the tomogram. Scale bars : 50 nm.

Figure 5A shows a central section across a representative cryo-tomogram containing E/EC1-2-liposomes interacting through numerous junctions. As signal to noise ratio is very low in cryo-EM images, the cryo-tomogram has been denoised by applying an anisotropic filter (Frangakis and Hegerl 2001).. To facilitate the visualization and interpretation, the contrast has been inversed. So, lipid membranes and E/EC1-2 fragments are in white. The lipid membranes appear as continuous white lines whereas E/EC1-2 fragments correspond to white elongated densities protruding out from lipid membranes. The tomogram section displayed in Figure 5A corresponds to the section where the inter-membrane distances of the two junctions noted 1 and 2 were minimal (see movie S1 in supplementary data). The insets (Figure 5 B-C and D-E) display a zoom of junctions 1 and 2 and a view of the same junctions in another tomogram section. Surprisingly, the molecular arrangements of the E/EC1-2 fragments in these two junctions were clearly different. Junctions from the inset 1 consisted of tangled and intertwined fragments forming a zigzag line (Figure 5B) whereas junctions represented in the inset 2 comprised straight, tilted zipper-like fragments similar to a ladder with an average repeated distance of 5.1 nm (Figure 5D). These molecular arrangements were conserved even when observed from different positions into the tomogram (Figure 5C and E). The inter-membrane distances of junctions 1 and 2 were of 12 nm and 9 nm respectively, arguing for different packing and consequently different E/EC1-2 adhesive interactions.

## DISCUSSION

This study aims at understanding how calcium ions impact the interplay properties of E/EC1-2, the two first E-cadherin extracellular modules, through observation in cryo-EM and cryo-ET of a very simple biomimetic system with minimal physical constraints consisting in E/EC1-2 decorated proteoliposomes. Previous observations showed that liposomes functionalized with E/EC1-5 or VE/EC1-4 cadherin fragments, the full-length or 4 outermost modules of E- and VE-cadherin respectively (Brasch et al., 2011; Lambert et al., 2005) form junctions in presence of calcium. As observed in this study, such junctions are also present with E/EC1-2 decorated liposomes in presence of calcium. It was reported that calcium binding to E/EC1-2 impedes the relative flexibility of EC1 vs EC2 modules (Häussinger et al., 2004; Pertz et al., 1999). As none junctions were observed with liposomes decorated with E/D103A fragments in presence or absence of 1 mM calcium, we can suppose that E/EC1-2 rigidification plays a role in the formation of *trans* interactions. Indeed, E/D103A is known to be 10 times less efficient than the WT protein (250 μM vs 25 μM) (Courjean et al., 2008), behaving as the apo-E/EC1-2 independently of the calcium conditions (Cailliez & Lavery, 2005) and therefore used as a non-adhesive control. Thus, calcium-dependent adhesive activity of the E/EC1-2 fragment, on surface of liposomes is impaired by the E/D103A mutation, in line with data showing that the full-length E-cadherin bearing only one substitution is unable to mediate contacts between cells (Prakasam, Chien, Maruthamuthu, & Leckband, 2006). Our data show the ability of cadherins to organize adhesive junctions after calcium adding is contained into the E/EC1-2 outermost segment, reinforcing the idea that these two outermost modules play a major role in junction formation. These results should be related to the role played by EC3 to EC5 modules in the full-length E-cadherin behavior. Indeed, deletion of these modules (Chappuis-Flament, Wong, Hicks, Kay, & Gumbiner, 2001) and mutagenesis experiments affecting different calcium binding pockets clearly reduce cadherin-mediated cell adhesion (Handschuh et al., 2001). Consequently, we can suppose that calcium-mediated EC1-5 arrangements may result from self-associated properties of the EC1-2 segment, and the glycosylation moieties on EC3 to 5 modules could play a regulatory role, by tending to disrupt *cis* interactions and ultimately destabilize or give plasticity to the contact structure.

Our experiments also revealed that calcium-triggered rigidification of E/EC1-2 fragments induced their nucleation and clustering giving rise to quasi-crystalline E/EC1-2 nanoplatforms. In other words, calcium-mediated E/EC1-2 structural rearrangements lead to lateral *cis* interactions. The cohesive state of these cadherin intermolecular *cis*-interactions had to be stable as these structural features could cover large areas of the surface of liposomes. Such *cis*-arrays have never been observed with E/D103A-liposomes in presence or absence of calcium. This is in contradiction with previous results indicating that *cis* interactions were not observed in NMR measurements of E/EC1-2 (Häussinger et al., 2004) or in single-molecule FRET experiments where E-cadherins were placed in close *cis* proximity (Zhang, Sivasankar, Nelson, & Chu, 2009). However, this is in agreement with recent work showing by single-molecule tracking of cadherin extracellular domains on supported lipid bilayers that revealed the formation of oligomers and *cis*-clusters in the absence of trans-interactions (Thompson et al., 2020, 2019). The design of experiments is certainly the most obvious explanation for these discrepancies. Indeed, the first experiments used E-cadherin fragments in solution while the seconds used E-cadherin fragments linked to a lipid membrane inducing structural constraints that do not exist or at least are reduced in solution. Moreover, for biochemical reasons E-cadherin fragments were produced for a long time with a methionine (Met; M) at the N-terminal end. In our hand, the behavior of M-E/EC1-2 and E/EC1-2 when attached to liposomes or lipid monolayers are different. Thus, we never obtained organized junctions when using M-E/EC1-2 fragments (Fig 6), suggesting that the impact of one amino-acid can be dramatic on the interaction of E/EC1-2 fragments. Nevertheless, the formation of junction with M-E/EC1-2 supposes *cis* interactions that were never observed from similar fragments in solution by NMR (Häussinger et al., 2004). Consequently, it suggests that interactions of E-cadherin fragments in solution or linked to a lipid membrane are extremely sensitive to the environment, explaining why literature on cadherins is sometimes contradictory.

**Figure 6:**
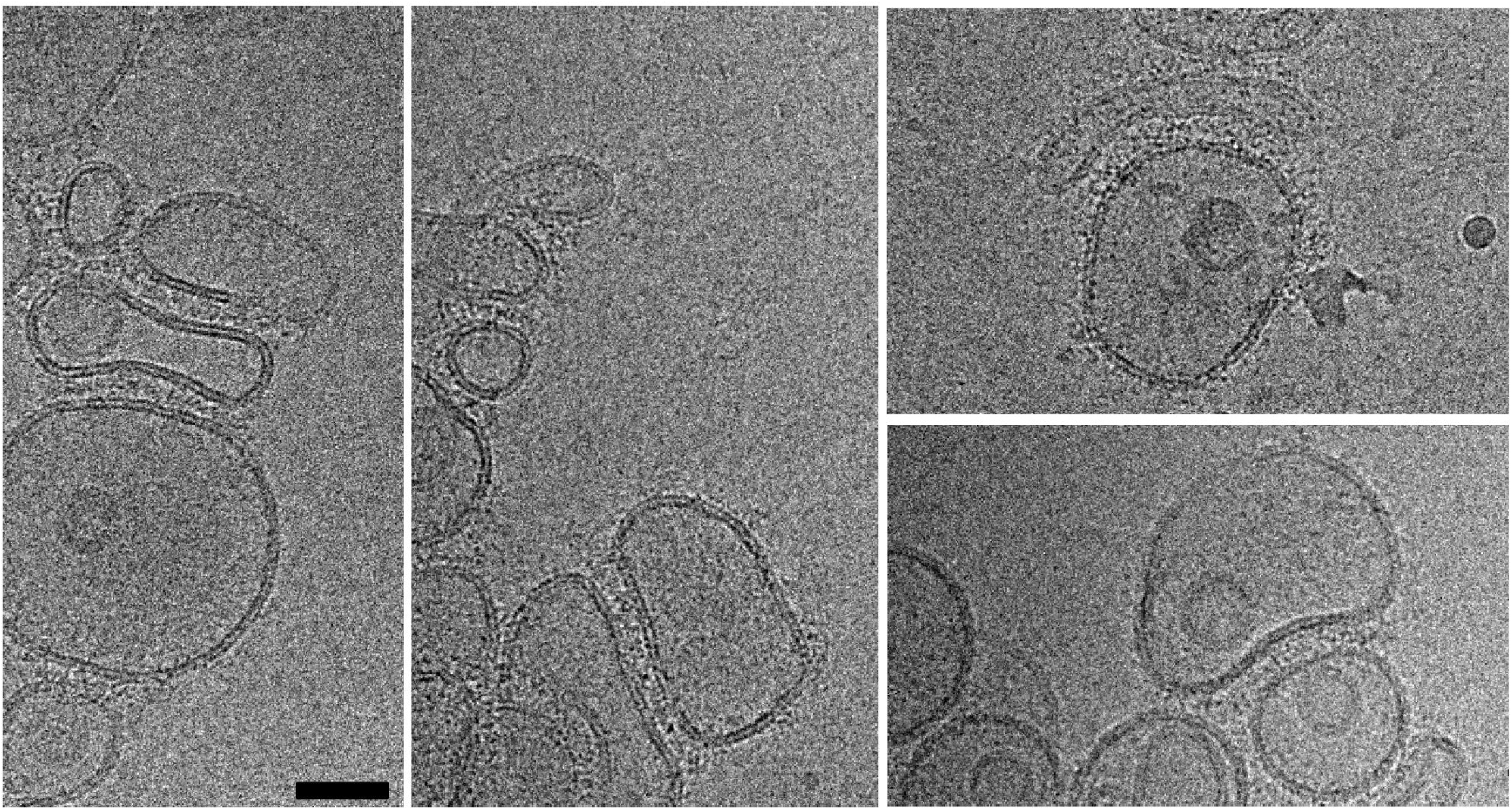
M-E/EC1-2 decorated liposomes incubated 10 minutes with 1 mM calcium. Some typical images observed with M-E/EC1-2 proteoliposomes in presence of 1 mM calcium (Ca^2+^). Liposomes made from DOPG/DOGS-Ni-NTA (4/1 : w/w) lipid mixture. Many junctions can be observed but in no apparent order. Scale bar : 50 nm.

As reported previously on mimetic system or from Human Embryonic Kidney (HEK293T) cells plasma membrane (Singh et al., 2017), one main effect of calcium binding on E-cadherin is to facilitate the spontaneous organization of cadherin *cis*-clusters, and our data suggest that this ability is also contained in E/EC1-2 modules. It is important to note it is the first time that *cis*-interactions in the absence of *trans*-interactions are directly visualized.

Surprisingly, in our experiments, we only observed E/EC1-2 *cis*-arrays in short timescale, ranging from 5 to 10 min after calcium addition; the maximal number of well-organized junctions being observed between 10 and 15 min. The regular nature of E/EC1-2 junctions indicates an identical pattern of E/EC1-2 interactions. Beyond 15 min, junctions became less ordered suggesting now the presence of different types of interaction between E/EC1-2 fragments. If we had the same probability to form *cis*-interaction between two free E/EC1-2 fragments and between a free E/EC1-2 fragment with one already involved in a junction, we would expect to regularly observe *cis*-arrays between 15 to 30 min, at least as long as some E/EC1-2 fragments are still free and not involved into junctions. Considering the fact that we never observed *cis*-arrays between 15 and 30 min and that *cis*-interactions are still present since leading to aggregation of E/EC1-2 decorated liposomes through reinforcement of junctions, we can suppose of a cooperative effect of *trans* interactions on *cis* interactions.

In summary, we showed that calcium first induces E/EC1-2 *cis* interactions allowing the formation of *cis*-arrays. These nanoplatforms, due to their high protein density, may subsequently facilitate and contribute to the rapid formation of stable junctions; it means the formation of *trans* S-dimers, which are known to have a low dissociation constant (*K_D_*) (Vendome et al., 2011). Thus, *cis* interplays favor the formation of *trans* interactions. Then, E/EC1-2 fragments are recruited at the level of junction, inducing the protein depletion of liposome surfaces. In other words, *trans* interactions favor the local recruitment of E/EC1-2 fragments through *cis* interactions. This cooperativity, *cis* to *trans* and *trans* to *cis*, deduced from our experiments is in agreement with observations carried out from E-cadherin decorated lipid droplets suggesting a *cis* and *trans* cooperativity in the formation of cadherin adhesion (Pontani, Jorjadze, & Brujic, 2016) and, from very recent work performed by super-resolution microscopy and intermolecular single-molecule FRET using a biomimetic lipid bilayer cell adhesion model, showing that *cis* and *trans* interactions are mutually cooperative (Thompson et al., 2021). Here again, our data suggest that *cis*/*trans* and *trans*/*cis* cooperativity of cadherin ectodomains is contained in E/EC1-2 modules.

Cryo-ET study of E/EC1-2 decorated liposomes allowed us to distinguish two types of junctions, presenting different E/EC1-2 molecular architectures: a zigzag-like and a ladder-like organization of junctions respectively (Figure 7A). As the maximum length of an E/EC1-2 fragment determined from atomic structures in the PDB database (1EDH) ranges from 8 to 9 nm, the junction displaying the ladder-like organization having an inter-membrane distance of about 9 nm, certainly represents head to tail interacting E/EC1-2 fragments, with the EC1 module of one fragment interplaying with the EC2 module of the opposite fragment (Figure 7B), as previously suggested and described in *(a)* EM studies of *adherens junctions* (Tariq et al., 2015), *(b)* X-ray crystallography (X-ray) (Boggon et al., 2002), *(c)* Biomembrane Force Probe (E. Perret, Leung, Feracci, & Evans, 2004) and *(d)* Surface Force Apparatus (S. Sivasankar, Brieher, Lavrik, Gumbiner, & Leckband, 1999) experiments or from X-ray structure of human P-cadherin EC1-2 (Dalle Vedove, Lucarelli, Nardone, Matino, & Parisini, 2015). The junction displaying the zigzag-like organization has an inter-membrane distance of 12 nm and the repetitive motif in Figure 5C is very similar to what we would expect with multiple interactions of E/EC1-2 fragments forming X-dimers (1EDH) (Figure 7C) (Nagar et al., 1996). This is the first time it is possible to observe simultaneously different structural organizations for cadherin fragments, from a biomimetic system where E/EC1-2 are linked to fluid lipid membranes, a system closer to reality than traditional X-ray or NMR experiments performed from soluble E-cadherin fragments. Another interesting point revealed from cryo-ET is that E/EC1-2 junctions are not perfectly ordered when moving through the width of junctions in tomogram, meaning that even in apparent well-ordered E/EC1-2 junctions, fragments can intrinsically interact through various conformations. In addition, the progressive disorder observed inside E/EC1-2 junctions in the second part of our calcium incubation experiments and related to the reinforcement of *cis* interplays, supports this idea. E/EC1-2 fragments when they are linked to a lipid membrane are dynamical modules that can simultaneously adopt various interaction states with neighboring fragments in junctions, a dynamical feature that has to be related to the diversity of E/EC1-2 interactions revealed by X-ray crystallography or NMR. If we summarize E/EC1-2 properties when they are attached to lipid membrane, E/EC1-2 fragments induce cadherin *cis*-clustering, the ability to form junctions, are involved in the mutual cooperative effect between *cis* and *trans* interactions, and can adopt various interaction states in junctions. These properties should be related to the fact that junctions visualized in cryo-EM with liposomes functionalized with E/EC1-5 (Harrison et al., 2011) (Figure 4 A-B) are not very-well structured, especially if we compare to the regular organization of desmosome (Al-Amoudi et al., 2011).

**Figure 7:**
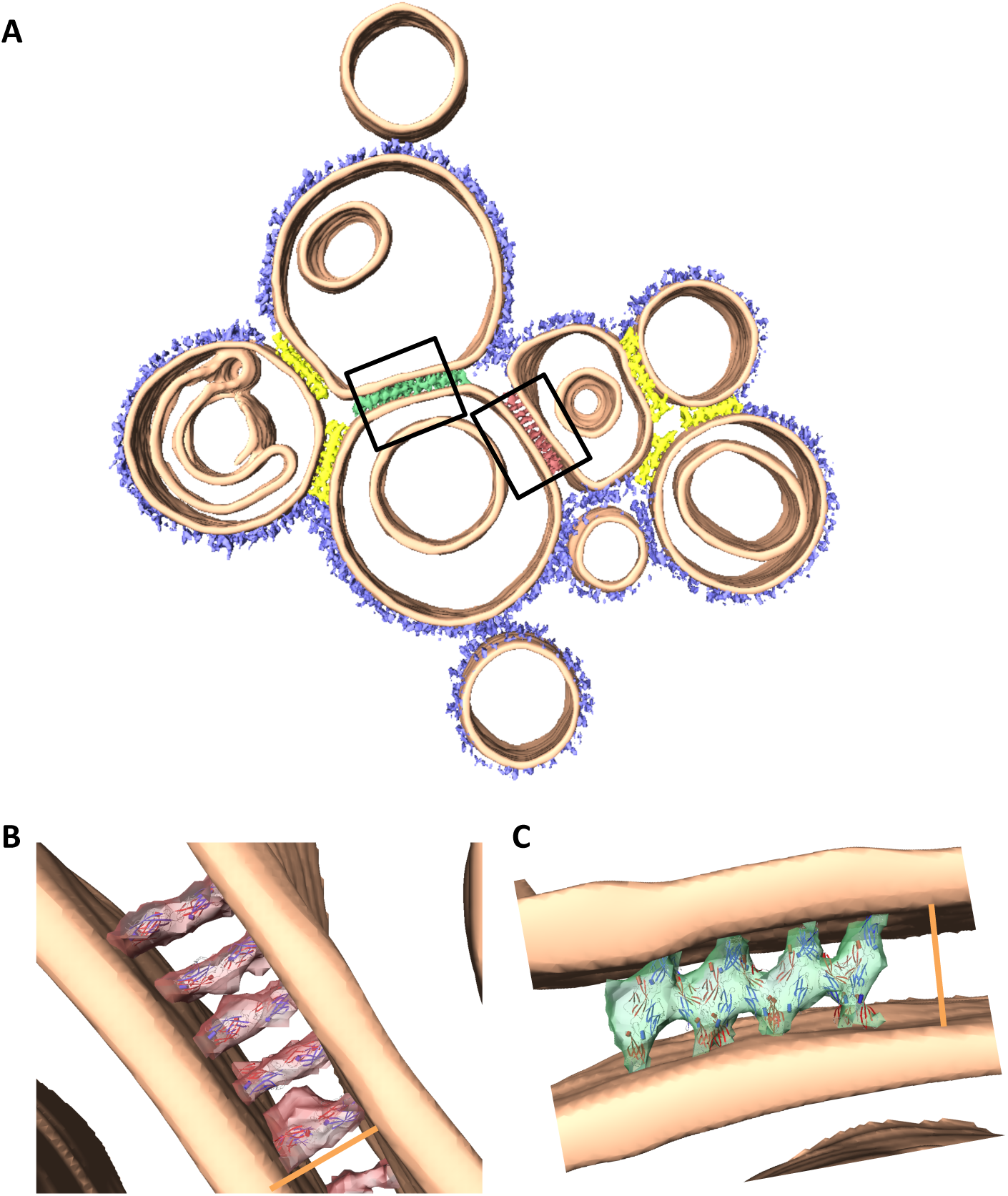
Models of molecular organization of E/EC1-2 junctions. **A** : Segmentation of the cryo-electron tomogram shown in Figure 5. **B and C** : Superimposition of models built using the atomic structure of E-cadherin domains 1 and 2 in complex with calcium, obtained by X-ray crystallography (PDB: 1EDH), with tomogram sections of (**B**) the ladder-like junctions and (**C**) the zigzag-like junctions shown in Figure 5C and 5E, respectively. The same scale bar (orange) has been added to visualize the size difference between the junctions.

In conclusion, this study provides new insights into the impact of calcium ions and suggests that through their intrinsic ability to form multiple calcium-dependent *cis* and *trans* interactions, E/EC1-2 modules play a key role in the formation of cadherin junctions contributing to the dynamics and plasticity of cadherin-mediated cell junctions.

## MATERIALS AND METHODS

### Protein production and purification

The construct to encode the first two extracellular domains fused to a C-terminal hexahistidine tag and the purification protocol were previously reported in Perret E et al. (Emilie Perret et al., 2002). Briefly, primers were used to amplified by PCR the first two extracellular domains E/EC1-2. The resulting fragment was inserted into a pET24a plasmid in order to fused a histidine tag at the C-terminal of the domains. For production of the domains EC1-2, a transformed colony picked from an agar plate was inoculated in 500 ml of Terrific Broth supplemented with 50 µg/ml kanamycin. The protein was expressed by the addition of isopropyl-ß-D-thiogalactopyranoside (IPTG) for 2 h. Cell pellets were resuspended in lysis buffer (4 M Urea, 50 mM Na_2_HPO_4_, pH 7.8, 20 mM imidazole and 20 mM ß-mercaptoethanol), and the lysate was then clarified by centrifugation, followed by purification using Ni^2+^-NTA-agarose resin (Qiagen). Purity and folding of the purified domains were finally verified using SDS-PAGE, western blotting, circular dichroism and mass spectrometry. It is important to note that comparing to previous work, in this study the N-terminal of the EC1-2 domains is free from additional aminoacids, such as a Methionine, as previously reported (Courjean et al., 2008). To avoid both dimer formation through S-S bonds and calcium contamination, all purified fragments were subject to reducing agents and calcium chelators.

### Preparation of E/EC1-2 proteoliposomes

1,2-dioleyl-sn-glycero-3-phospho-(1’-rac-glycerol) (sodium salt) (DOPG) and 1,2-dioleyl-sn-glycero-3-[(N-(5-amino-1-carboxypentyl)iminodiacetic acid)succinyl] (nickel salt) (DOGS-Ni-NTA) were purchased at Avanti Polar Lipids, Inc. The lipid mixture DOPG/DOGS-Ni-NTA (4/1 : w/w) in chloroform/methanol (9/1 : v/v) were prepared. The chloroform was evaporated under a stream of nitrogen and the lipids film was resuspended in 1 mL of 20 mM Tris-HCl, 150 mM NaCl and 0.8 µM β-mercaptoethanol pH 8 buffer. Lipids mixtures were homogenized by three cycles of freeze-thawing. 250 µL of this lipids solution was extruded through a polycarbonate membrane of 0,1 µm. The liposomes solution had a final concentration of ∼2 mg/mL and was stored at 4 °C under nitrogen. The liposomes (DOPG/DOGS-Ni-NTA : 4/1) and proteins (E/EC1-2 or E/D103A) solutions were mixed respectively in a final concentration of 250 µg/mL and 100 µg/mL. The mixtured were incubated 1 hour before addition of calcium at a final concentration of 1 mM. In each experiment, the preparation of cryo-EM grids was carried out 5, 10, 15 and 30 min after calcium incubation.

### Cryo-electron microscopy

Three microliters of liposomes or proteoliposomes were applied to glow discharged Quantifoil R 2/2 grids (Quantifoil Micro Tools GmbH, Jena, Germany) or Lacey grid (Ted Pella inc.), blotted for 1s and then flash frozen in liquid ethane using a CP3 cryo-plunge (Gatan inc.). Before freezing, the humidity rate was stabilized at about 95 %. Cryo-EM was carried out on a JEOL 2200FS FEG operating at 200 kV under low-dose conditions (total dose of 20 electrons/Å2) in the zero-energy-loss mode with a slit width of 20 eV. Images were taken at a nominal magnification of 50,000X corresponding to a calibrated magnification of 45,591X with defocus ranging from 1.4 to 2.5 µm.

### Cryo-electron tomography

Samples were prepared as described for cryo-electron microscopy, except that sample concentration was increased by a 4-fold factor and treated with 5 nm colloidal gold fiducial markers (BBI International, Agar Scientific) before sample application.

The tomograms were computed from tilt series collected on a 4k X 4k slow-scan CCD camera (Gatan inc.) from -60° to 60° using a JEOL 2200FS microscope operating at 200 kV at approximately 5 µm underfocus with a nominal magnification of 30,000X. The acquisition was performed semi-automatically using a version of SerialEM (http://bio3d.colodaro.edu) adapted for operating JEOL microscope allowing recording of individual images in the zero-energy-loss mode with a slit width of 20 eV. The total dose for one tilted serie is estimated to 90 e^-^/Å^2^. The resulting pixel size corresponds to 0.4 nm on the specimen.

### Image analysis

Image stacks were aligned either manually with the eTomo suite of programs, or automatically using the software RAPTOR (Amat et al., 2008). Tomographic reconstructions were constructed using the eTomo suite of programs and visualized using the 3dmod software package (Kremer, Mastronarde, & McIntosh, 1996). Cryo-electron tomograms were denoised by applying an anisotropic filter (Frangakis & Hegerl, 2001). The segmentation of cryo-electron tomogram was carried out using the AMIRA software (Thermo Fisher Scientific).

## Supporting information

Movie S1: visualization of sections through the cryo-electron tomogram used in Figure 5.

## ACKNOWLEDGEMENTS

We would like to thank the « Unité mixte de services » Biocampus (Montpellier) and the CBS for their constant support. The CBS is a member of the French Infrastructure for Integrated Structural Biology (FRISBI), a national infrastructure supported by the French National Research Agency (ANR-10-INBS-05).

## AUTHOR CONTRIBUTION

Joséphine Lai-Kee-Him, Aurélien Fouillen, Matthew Chee, Nicolas Lemercier, Hélène Feracci and Patrick Bron participated in the design of the experiments and in interpretation of results. Joséphine Lai-Kee-Him performed most of the experimental work and data analysis. Nicolas Lemercier performed cryo-EM observation of M-E/EC1-2 decorated liposomes. Aurélien Fouillen carried out the tomogram reconstructions and Matthew Chee performed the segmentation of cryo-electron tomograms. Aurélien Fouillen and Patrick Bron wrote the manuscript. All authors discussed the results and manuscript. Patrick Bron provided scientific and financial support, and supervised this project.

## ABBREVIATIONS

2D: two-dimensional
3D: three-dimensional
Å: Angström (10^-10^ meter)
µM: micro-molars (10^-6^ M)
A: Ala = Alanine
Ca^2+^: Calcium ions
Cryo-EM: Electron microscopy in cryogenic conditions
Cryo-ET: Electron tomography in cryogenic conditions
D: Asp = Aspartic acid
DGS: 1,2-dioleyl-sn-glycero-3-[(N-(5-amino-1-carboxypentyl)iminodiacetic acid)succinyl]
DOPG: 1,2-dioleyl-sn-glycero-3-phospho-(1’-rac-glycerol)
E: Glu = Glutamic acid
EC: E-cadherin module
E/EC: Extracellular E-cadherin module
EM: Electron microscopy
eV: Electron voltage
FEG: Field emission gun
FRET: Forster resonance energy transfer
FT: Fourier transform
IPTG: isopropyl-ß-D-thiogalactopyranoside
K: Lys = Lysine
KD: Dissociation constant
kV: Voltage en kilo (1000 V)
M: Methionine
min: minutes
mM: milli-molar (10^-3^ M)
NMR: Nuclear magnetic resonance
Ni-NTA: Nickel-nitrilotriacetic acid
nm: nano-meters (10^-9^ m)
SDS-PAGE: Sodium dodecyl sulfate polyacrylamide gel electrophoresis
sec: seconds
WT: Wild-type
X-rays: X-ray crystallography

## Supplementary Data

**Movie S1:** visualization of sections through the cryo-electron tomogram used in Figure 5.

